# Understanding the host-microbe interactions using metabolic modeling

**DOI:** 10.1101/2020.06.12.147918

**Authors:** Jack Jansma, Sahar El Aidy

**Affiliations:** Department of Molecular Immunology and Microbiology, Groningen Biomolecular Sciences and Biotechnology Institute (GBB), University of Groningen, Groningen, Nijenborgh 7, 9747 AG Groningen, The Netherlands

**Author notes:** **Corresponding author** Sahar El Aidy. Department of Molecular Immunology and Microbiology, Groningen Biomolecular Sciences and Biotechnology Institute, University of Groningen, Nijenborgh 7, 9747 AG Groningen, The Netherlands., E. **E mail addresses:** Jack Jansma < >, Sahar El Aidy < >.

**Keywords:** Flux balance analysis, Gut microbiota, Probiotics, Metabolic model, Microbial community

## Abstract

The human gut harbors an enormous number of symbiotic microbes, which is vital for human health. However, interactions within the complex microbiota community and between the microbiota and its host are challenging to elucidate, limiting development in the treatment for a variety of diseases associated with microbiota dysbiosis. Using *In silico* simulation methods based on flux balance analysis, those interactions can be better investigated. Flux balance analysis uses an annotated genome-scale reconstruction of a metabolic network to determine the distribution of metabolic fluxes that represent the complete metabolism of a bacterium in a certain metabolic environment such as the gut. Simulation of a set of bacterial species in a shared metabolic environment can enable the study of the effect of numerous perturbations, such as dietary changes or addition of a probiotic species in a personalized manner. This review aims to introduce these applications of flux balance analysis to experimental biologists and discusses its potential use to improve human health.

## Metabolic networks

The gut microbiota is the community of microorganisms residing in the gut and include commensal, symbiotic, and pathogenic bacteria. Under normal circumstances, the gut microbiota and the host are in symbiosis [1]. Disruption of the symbiosis is detrimental for host health and can result in disease including gastrointestinal disorders such as inflammatory bowel disease [2], metabolic disorders such as diabetes mellitus [3], and mental disorders such as autism spectrum disorder [4], and major depressive disorder [5]. To understand the symbiosis, the different members of the gut microbiota, and the way they communicate with each other and with the host need to be known. The microbiota communicates via production of metabolites [6]. Therefore, it is key in the field of host-microbe interactions to identify which microbial members are present and what their metabolic output is. However, this does not fully elucidate the dynamic interactions within the microbiota, and between host and the microbiota, since the metabolic output of microorganisms is dependent on their surroundings [7]. Therefore the metabolic output and, in turn, the interactions between host and microbiota is different among individuals [8], making successful treatment of the aforementioned disorders more challenging. Although experimental approaches are crucial to the progress of the microbiota field, they are not able to fully capture the mechanisms, interactions, and behavior due to the huge complexity of the environment in the gut. These limitations have led to the development of a complimentary approach to completely understand the relationship between host and microbe; bacterial metabolic networks [9]. In this approach, bacterial interactions can be visualized in the form of a metabolic network. The bacteria, metabolites, and the host cells comprise the nodes of the metabolic network. Biological processes such as conversions, uptake, and secretion are represented by the edges [10] (Figure 1).

**Figure 1:**
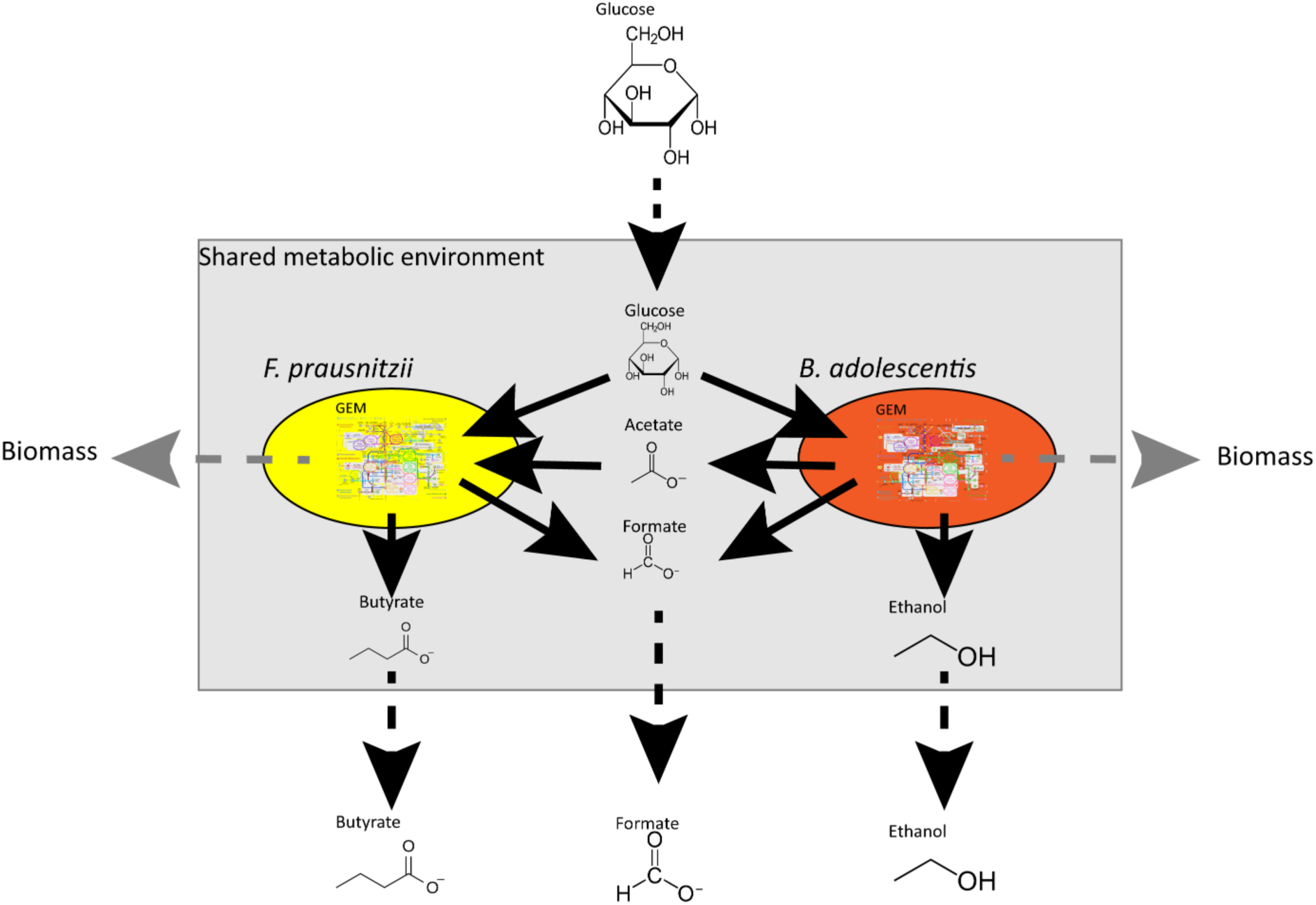
A simplified metabolic network of a bacterial community consisting of *Faecalibacterium prausntizii* and *Bifidobacterium adolescentis* (adapted after El-Semman et al. [2]). Fluxes are represented by arrows. Solid black arrows indicate uptake, and secretion reactions of the bacteria, dashed black arrows indicate the flow of metabolites in or out of the system, and dashed grey arrows indicate the formation of new biomass, where metabolites cannot be secreted by the bacteria anymore, thus leaving the system.

Metabolic network models can be used to predict the effect of alterations in the metabolic network *in silico*. A constraint-based reconstruction and analysis (COBRA) approach is often used to simulate the operation of a metabolic network under different external nutrient conditions. A COBRA method suitable for investigating the metabolism of microbiota is flux balance analysis (FBA)[11]. A flux is the rate of turnover of a metabolite through a metabolic pathway. To perform FBA, all the fluxes in the network should be represented by a set of linear equations. The equations are placed in a stoichiometric matrix, which consists of the substrates, products, and directionality of the reactions (Figure 2). Next, FBA uses constraints to limit the flow of metabolites through the network and calculates the distribution of metabolic fluxes in the metabolic network for a given objective function (OF), resulting in an optimal distribution of fluxes (Figure 3). For example, an OF that maximizes the production of the short chain fatty acid (SCFA) acetate will result in a different flux distribution compared to an OF that maximizes butyrate production. Other examples of OFs include minimizing the production of a particular metabolite, maximizing cell growth or for a community of bacteria maximizing growth of a single species or maximizing the growth of the whole community [11]. FBA works at steady-state; i.e. the amount of the metabolite produced is equal to the amount of the metabolite consumed. Accordingly, the set of linear equations is formulated as:

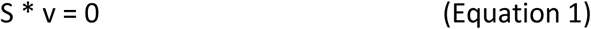

**Figure 2.**
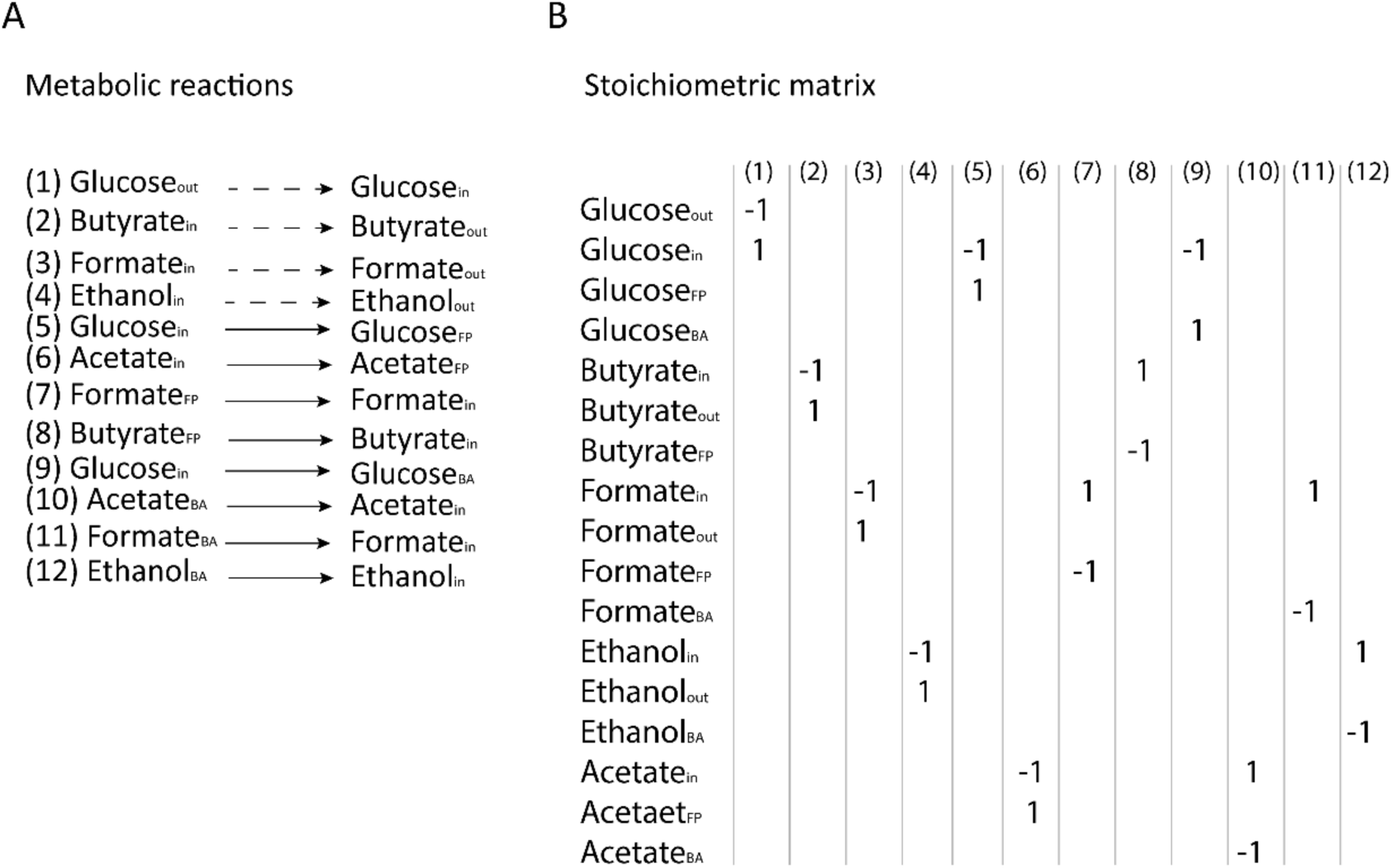
The metabolic reactions indicated in Figure 1 and their stoichiometry. (A) The metabolic reactions depicted in Figure 1. Dashed arrows indicate reactions between the outside of the system and the shared metabolic environment. Solid reaction arrows indicate reactions between the shared metabolic environment and the bacteria, (B) Stoichiometric matrix of the reactions depicted in panel A. BA is *B. adolescentis* and FP is *F. prausnitzii*.

**Figure 3:**
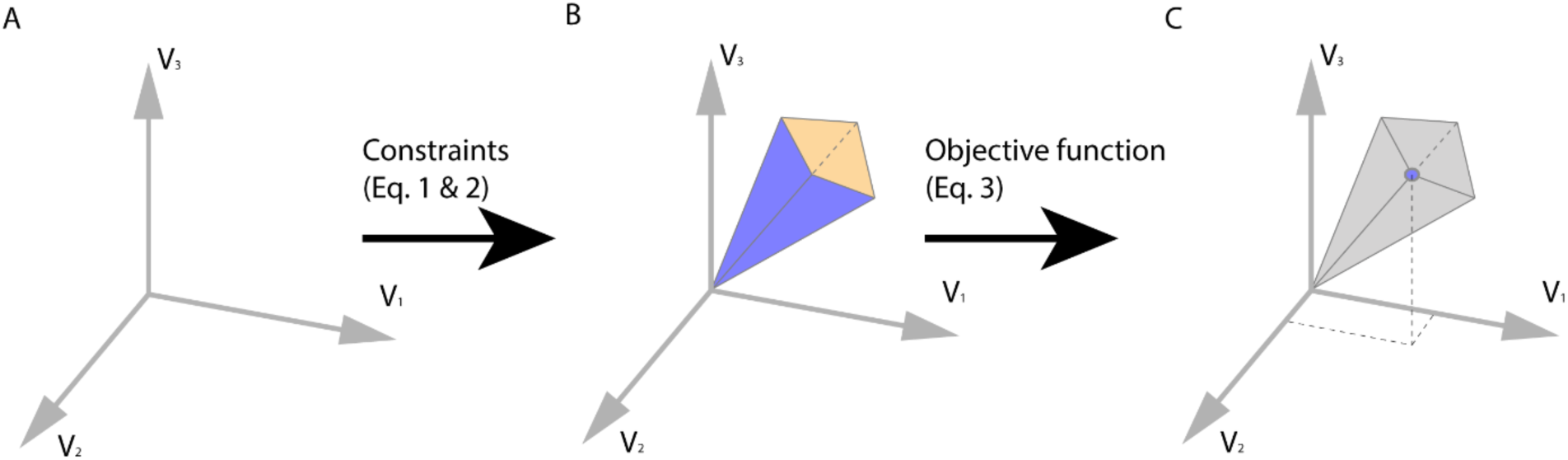
Visual representation of the concept of FBA. (A) A solution space of the flux distribution in a system. (B) The allowable solution space is the result of addition of constraints depicted in equations 1, and 2. The equations limit the available flux distributions. (C) Addition of an objective function (equation 3) determines the optimal solution for the flux distribution in the system. *Adapted after Orth et al*. [3].

Where (S) represents a stoichiometric matrix, and (v) represents the flux distribution. The formula describing a biomass function contains all the metabolites in the system that are required to build a new cell. In other words, a biomass function simulates cell growth. Importantly, since ***Equation 1*** depicts a set of linear equations and there are typically more reactions than compounds, there is more than one flux distribution possible. Constraints are imposed on a flux as upper- and lower- bounds to limit the maximum and minimum values that each flux can take. Constrains could reflect media conditions, where uptake and secretions rates are limited, or reaction velocities of internal enzymes obtained from experimental data [12, 13]. Accordingly, each flux with constraints is formulated as:

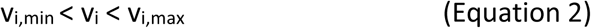

Similarly, the OF is formulated as:

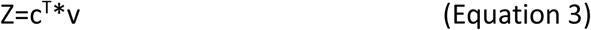

where (Z) is the solution of the OF, (C) is the vector of weights, indicating how much each reaction contributes to the OF and (T) represents the matrix transpose. For example, when a single flux is maximized or minimized, C is a vector of 0s with a single 1 [14, 15]. However, formulating an OF can be challenging and is fully dependent on the research question [16, 17]. FBA is a versatile tool to employ for many purposes. By adjusting the upper- and lower-bounds of metabolites, growth on different media [18] or, in the case of the gut microbiota, changes in diet, can be simulated. By setting the flux of a certain metabolite to zero, a gene knock-out or the absence of a member of the microbiota can be simulated [19, 15, 11]. In this way, the viability of a microbial community under different conditions as well as the effect of adding new species to a bacterial community on host health can be estimated [16]. For example: FBA used with a metabolic model of *Bifidobacterium adolescentis* shows a reduction in the production of formate, ethanol and acetate as well as a reduction in biomass formation if lactate production is increased. The OF in this example is maximizing biomass production (Figure 4) showing that the flux distribution changes with altering the environment [20]. To investigate a microbial community rather than a single bacterium (Figure 4), another bacterium, *Faecalibacterium prausnitzii*, is placed in the same metabolic environment as *B. adolescentis. F. prausnitzii* needs acetate to grow well on glucose and to produce butyrate [21]. Since acetate is not supplied, *F. prausnitzii* can use the acetate produced by *B. adolescentis* to grow and to produce butyrate. Altering the ratio between both bacteria results in an increase in butyrate production when more *F. prausntizii* is placed in the system compared to a situation with more *B. adolescentis*. The OF in this example is minimization of glucose consumption and the total biomass formation is kept constant (Figure 5). This example shows that FBA and metabolic networks can be used to investigate interactions between bacteria.

**Figure 4:**
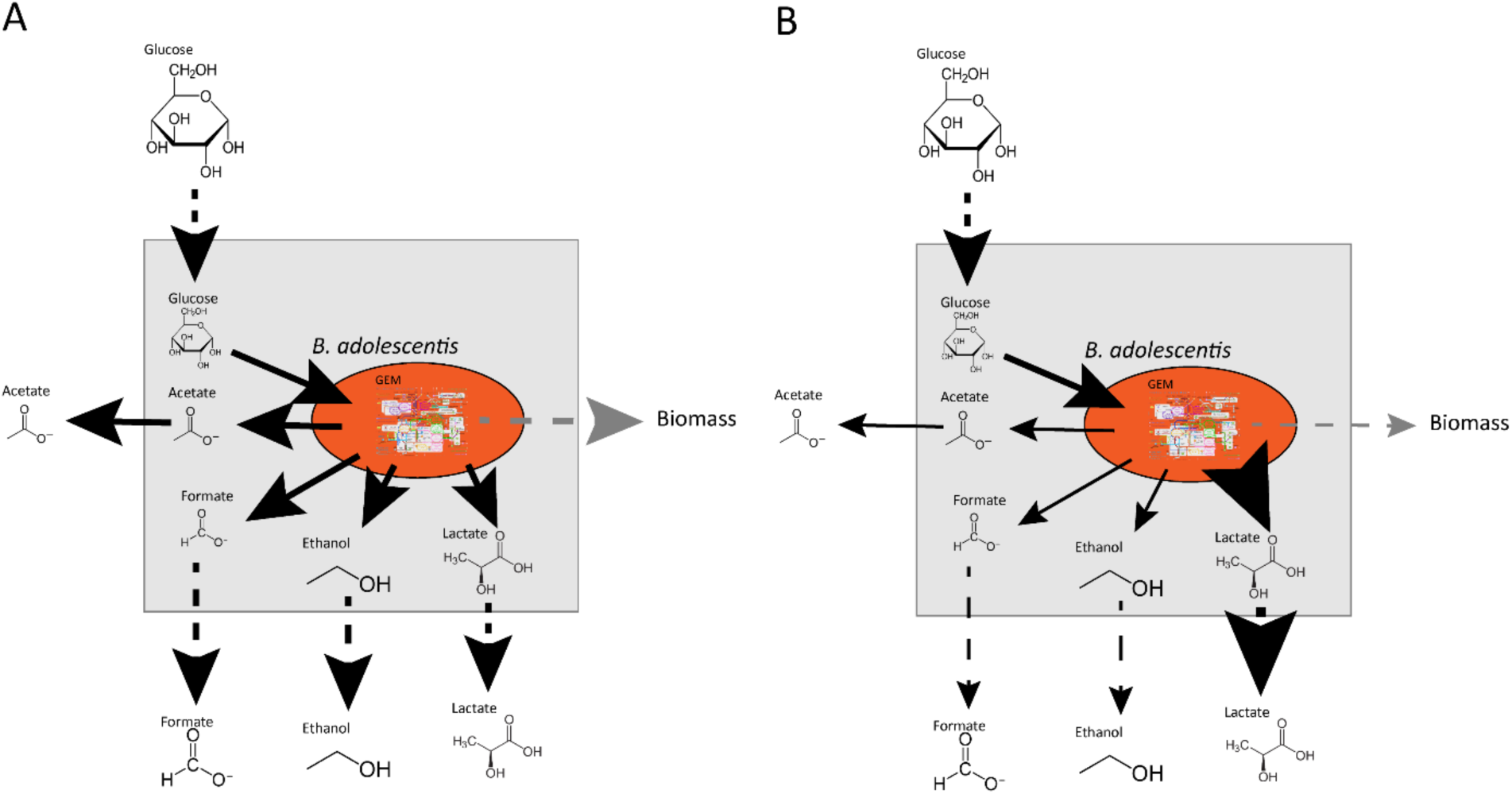
An example of the use of FBA in metabolic modeling of an organism. The OF panels A and B is maximizing the production of the biomass. The thickness of the arrows indicate the amount of flux, where a thicker arrow indicates a higher flux. When the production of lactate is manually altered in panel B, the flux distribution changes, whereby the flux of acetate, formate, ethanol and biomass is lowered compared to panel A. *Adapted after El-Semman et al*. [2].

**Figure 5.**
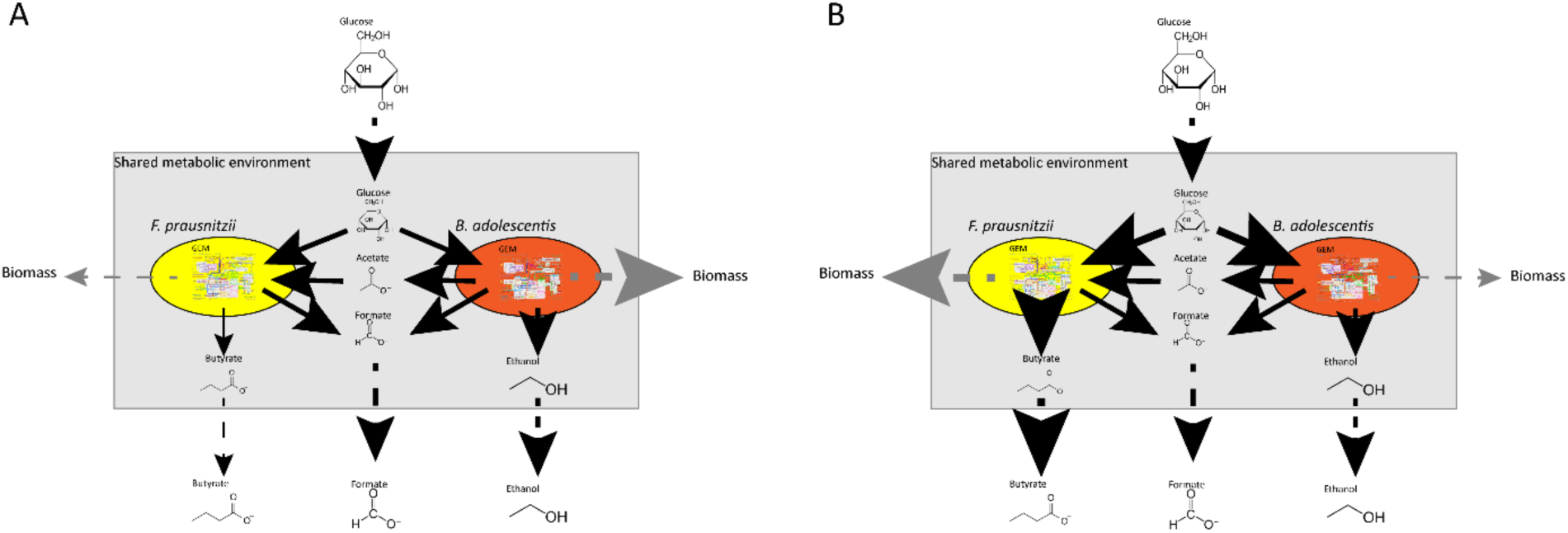
An example of the use of FBA in the modeling of a bacterial community consisting of two gut bacteria: *F. prausnitzii* and *B. adolescentis*. Fluxes are represented by arrows. Solid black arrows indicate uptake and secretion reactions of the bacteria, dashed black arrows indicate the flow of metabolites in and out of the system and dashed grey arrows indicate the formation of new biomass, where metabolites can no longer be secreted by the bacteria, thus leaving the system. The amount of flux is represented by the thickness of the arrows, a higher flux is a thicker arrow. The compositions is determined by the amount of biomass produced. The total biomass in this example is the same. The OF is the minimization of glucose uptake. *Adapted after El-Semman et al*. [2].

The examples depicted in figures 4 and 5 show the usefulness of metabolic models in studying metabolic capabilities of bacteria, and interactions among bacteria. However, since FBA works at steady-state, it gives only one optimal distribution of fluxes for a given OF. This does not capture dynamic changes in metabolite levels and dynamic interactions between bacteria. Dynamic changes are an important factor when studying bacterial communities, because cells are dividing and dying. Additionally, in the human gut there is movement of cells from the upper part of the intestine to the lower part due to gut motility and metabolites levels change dynamically during the day due to the cycles of food intake [22, 23, 24]. Thus, capturing dynamic changes in metabolic networks is necessary for studying the gut microbiota. To capture dynamic changes, dynamic FBA (DFBA) has been developed. DFBA uses ordinary differential equations (ODE) to couple FBA to a kinetic model [25]. This can be done using three approaches: static optimization approach (SOA), dynamic optimization approach, or direct approach [26]. The most widely used approach is SOA. Essentially, SOA makes a series of snapshots using FBA. The starting conditions of each snapshot is determined by the outcome of the previous snapshot [27]. For example: Mahadevan et al. used the SOA approach to investigate the restructuring of the metabolic network of *Escherichia coli* during a diauxic shift. The authors simulated a batch culture of 10 h, which was divided into 10.000-time intervals. The concentration of the metabolites at the start of each interval was directly calculated from the previous interval. This approach showed that during the first 4.6 h the oxygen, and glucose uptake rates of the cells were the limiting constraints on biomass formation. During the second phase between 4.6 h and 6.9 h the limiting constraint on biomass formation was the concentration of oxygen in the environment. During the last phase, where acetate is utilized from 6.9 h to 10 h the mass transfer coefficient is the limiting constrain on biomass formation. The computational values are similar to experimental values indicating the usefulness of DFBA to determine the limiting factors of growth in bacteria [28]. DFBA has been used to study communities of microorganisms including a community of *E. coli* and *Saccharomyces cerevisiae* [29]. However, DFBA can only be used in small communities of bacteria because adding more species in a network increases the amount of reactions dramatically, which consequently increases the time and costs for simulations. DFBA in microbial research is extensively reviewed elsewhere [26, 30]. To simulate microbial communities with FBA, not only the uptake and secretion reactions are necessary, but all the molecular capabilities of each bacterium in the metabolic network should be known. Thus, a metabolic network of each individual bacterium is needed. This is done by generating metabolic models of each bacterium directly from their sequenced genome.

## Genomic-scale metabolic models

To generate metabolic models, the metabolic capacity of the bacteria should be known and translated into a model that can be used by FBA. Genomic-scale metabolic models (GEMs), also known as genomic-scale metabolic reconstructions (GENREs), consist of all the metabolic capacities of a bacterium [31], and are automatically constructed from annotated genomes [32]. Several tools are available to construct GEMs, which are reviewed elsewhere [33]. However, not all genes of an organism are active during each growth phase, or in every environment [34, 35], hence, these automatically generated GEMs should be manually refined [36]. Recently, AGORA, a semi-automated GEM database of 818 members of the gut microbiota became available [37]. However, the AGORA GEMs differ from other available GEM databases such as BiGG, KBase, and CarveMe [38, 39, 40, 41]. Taxonomically, the AGORA database contains a wider variety of organisms and the GEMs are more generally constructed. This means that the user should add constrains dependent on the research question. Furthermore, the AGORA database is built to simulate the gut microbiota community in general. Therefore, the AGORA models have, for example, more carbon-uptake reactions compared to the BiGG database, which has a more limited number of bacteria and is generally used to simulate metabolic capabilities of a single bacterium and predicts possible changes in the metabolic capacity of this bacterium in case it harbors inoperative genes [39]. For example, the BiGG database contains numerous GEMs of *E. coli*, which is a heavily researched organism, thus the metabolic reactions and constraints can be accurately determined from literature. On the contrary, the AGORA database contains numerous under-investigated organisms, which results in less accurate GEMs. Therefore, GEMs from the AGORA database should be carefully looked at and proper constraints should be added before employing this database for simulation. Nevertheless, community efforts are being taken to standardize GEMs and assure the quality of the models [42]. To optimize GEMs of bacteria Kuang et al. combined FBA with an untargeted mass spectrometry-based approach to identify metabolites produced by *Citrobacter sedlakii*, a non-pathogenic bacterium found in human stool. The authors investigated the metabolic output of *C. sedlakii* by taking samples at different growth stages and analyzing the extracts using two liquid chromatography mass spectrometry (LCMS) approaches, reverse phase (RP) and hydrophilic interaction liquid chromatography (HILIC). The obtained data was compared to a predicted list of metabolites generated via FBA using a tool called MS_FBA. The comparison showed that the metabolic output of the GEM of *C. sedlakii* did not cover all the metabolites measured with LCMS, highlighting the need to determine the accuracy of bacterial GEMs and more precise genome annotation [43]. Lastly, when comparing flux distribution between different sets of microbial profiles, for example between patients and healthy controls, the metabolic environment of the microbial community should be known. This can be partly inferred from the diet of the patients, but never to a full extent. Another approach to infer the metabolic environment of the microbial community is to infer the metabolic environment from abundance distribution in individual samples. The Metabolic Analysis of Metagenomes using FBA and Optimization (MAMBO) approach combines 16S sequencing data from individual fecal samples with GEMs obtained from reference genomes of the sequenced bacteria to infer the metabolome in the sequenced fecal sample. The rationale behind this approach is that the metabolic environment causes the abundance profile in a given sample, because the metabolic potential of the bacteria indicate which species will thrive in the metabolic environment [44]. In other words, which metabolic environment fits the abundance profile best. To answer this question the authors placed GEMs of the bacteria in the same metabolic environment in a similar way as seen in Figure 5. Next the authors defined the starting concentration of the metabolites in the metabolic environment at random and performed FBA whereby the OF is defined as: the biomass functions of all the GEMs should be as close to the obtained abundance distribution. In a next step the metabolites in the metabolic environment are slightly altered and FBA with the same OF is performed again. If the Pearson correlation between the biomass functions and the abundance profile is higher compared to the previous metabolic environment the metabolite change is accepted and the process is repeated until the Pearson correlation does not improve Comparing the biomass distribution in the FBA results with the abundance profile obtained from the fecal sample shows which metabolic environment fits the abundance distribution best [45]. With the metabolic environment, microbial composition and metabolic capabilities known, the relationship between host and microbe can be studied.

## Metabolic networks in the gut

To understand the effect of the microbiota on the host, microbial metabolic networks can be extrapolated to include metabolic networks of the host. This enables predicting the effects of gut microbiota on the host and suggesting possible interventions to promote host health. Diener et al. constructed microbial metabolic networks from metagenome data of a cohort of Swedish people with and without diabetes mellitus and used FBA to analyze SCFA production. In this study, 16S abundance data was integrated with metabolic fluxes to construct personalized predictions of metabolic output and interventions such as dietary changes and medical treatment. The authors found a minimal overlap of resource utilization between microbes in different niches, suggesting an upper bound on alpha diversity in the gut. Next to ecological insights, the authors concluded from their model that SCFA production is highly specific per individual. Nonetheless production of butyrate and propionate was reduced in diabetic subjects compared to healthy controls and the overall SCFA production profile could be restored upon metformin treatment [19]. The authors did not measure metabolite levels *in vitro*, but the outcome is in line with other research [46]. Another study integrated publicly available microbial abundance data of healthy individuals and patients with inflammatory bowel disorders (IBD) in a metabolic network to study bile acids metabolism. The study showed that metabolism of bile acids needs to be done in a microbial community since individual bacteria cannot produce every bile acid. Furthermore, the study showed that the production of bile acids was lower in patients with IBD [47], which is in line with *in vivo* results [48]. However, the authors did not perform interventions such as dietary change or introduction of other bacteria in the metabolic network to predict possible treatment of IBD patients. Such interventions were taken in consideration in another study that used 16S sequencing data to construct metabolic networks of 28 Crohn’s disease (CD) patients, and 26 healthy controls. The study showed a higher production of SCFAs in the controls group. Furthermore, the authors increased the SCFAs produced in each of the constructed metabolic networks by altering the diet *in silico* and showed that dietary intervention can indeed have different results in each individual based on their microbiota composition [49], which is in line with experimental results [50].

Currently, several tools are available for modeling bacterial communities, such as OptCom, BacArena, MiMoSa, COMETS, FLYCOP and MICOM [19, 51, 52, 53, 54, 55, 56]. To fully capture the effect of the microbial community on the host, metabolic networks of the gut community need to be extended with GEMs of host cells. A number of GEMs for human tissue such as the liver, blood vessels, and gut epithelial cells are already available to expand the existing metabolic networks of the gut microbiota to include human cells [57]. Combining metabolic networks based on 16s sequencing data with a representative GEM of human metabolism may be used to investigate the effect of the microbiota metabolic output on host health. A representative GEM of human metabolism is Recon3D, which contains over 13,000 metabolic reactions and can be used to integrate host and microbiota metabolism [58, 59]. However, combining host GEMs and bacterial GEMs can be challenging due to the formulation of the OF and spatial organization. For example, the intestine is spatially organized where intestinal cells surround the microbiota. Accordingly, gut microbiota metabolites have the highest concentration in the lumen and will be first available for the microbiota and not the host cells [60, 61]. When simulating interactions between gut microbiota and the host, metabolic networks should take these gradients into account. Similarly, metabolic and pH gradients exist along the length of intestinal tract [62, 63] and the microbiota composition in the small intestine differs from that in the colon. Furthermore, bacterial cells travel from the upper part of the intestine to the colon to be ultimately shed in the feces [64]. One way to simulate these gradients and spatial organization is through the addition of empty compartments between the compartments of the host cells and the bacterial compartments in a metabolic model. The compartments are organized in a two-dimensional grid and each bacteria has its own compartment from which it can exchange nutrients. The compartments are connected through exchange reactions, thus bacteria can interact with each other. By adding compartments without bacteria in between compartments filled with bacteria, a new bacterium can fill the empty compartment, which can simulate movement or replication of the bacteria (figure 7). Furthermore the empty compartments simulate gradients, since not all the nutrients entering the empty compartment will move to the next compartment, which is filled with a bacterium [51]. Another way to simulate gradients and movement of bacteria is by employing differential equations. Van Hoek et al. used a metabolic model in which spatial organization is included to show that cross feeding can be an emergent property. In turn, cross feeding resulted in more microbial and metabolic diversity in the metabolic model. When diarrhea was simulated by increasing the flux speed through the system, the microbial diversity was destroyed [65]. Chan et al. used a similar approach, to show that aerobic and anaerobic bacteria separate into different niches based on the oxygen gradient [66]. These examples show the importance of the gut environment and these factors should be taken into account. Persi et al. integrated pH dependent activity of enzymes obtained from experimental data in the BRENDA database in a GEM of cancer cells. The authors showed that cancer cells proliferate differently *in silico* based on the pH [67]. However, integrating pH dependent activity in a GEM is heavily dependent on experimental work done *in vitro*. Since most enzymes from gut bacteria are under investigated, integrating pH dependent activity in GEMs used for metabolic modeling is currently not feasible.

Another problem in combining bacterial GEMs and host GEMs is formulating an appropriate OF. In the majority of metabolic models, the total biomass growth, or metabolic output is optimized. In the gut, the host benefits the most from a balance between the metabolic output of all bacteria, which requires a balance among the distribution of the different microbial species. To address this problem OptCom was developed. This approach uses two layers of OFs. The first layer maximizes the biomass formation of each individual species. The second layer maximizes the growth of the whole community, resulting in a more realistic growth distribution in a bacterial community [52]. The extension d-OptCom can be used in DFBA [1]. However, OptCom cannot easily be used with communities consisting of a large number of bacteria due to increasing computational time needed with increasing community complexity. Therefore, community and systems level interactive optimization (CASINO) was developed. The CASINO framework uses two layers of OFs, but differs from OptCom by optimizing both layers of OFs iteratively [69]. A problem with OptCom and CASINO is the description of the biomass function. As mentioned above, the host benefits the most from a balance in the metabolic output of all bacteria. Thus, for a healthy microbiota the community needs to function at steady-state. However each individual bacterium grows with a specific growth rate [70]. OptCom and CASINO do take bacterial growth into account, but not the community steady state. Therefore, optimizing biomass formation results in the domination of the fastest growing bacterium in the system, resulting in a flux distribution that does not represent the flux distribution in the gut community. To overcome this problem, SteadyCom was developed which takes the community steady-state into account [71]. So far, studies have been able to combine both host and bacterial cells in simplified metabolic models. Heinken et al. combined GEMs of *Bacteroides thetaiotamicron* and a generalized mouse cell [72] in a metabolic network. They showed that the presence of *B. thetaiotamicron* could influence the growth of the generalized mouse cell and metabolic dependency between the bacterial and host cells [73]. This metabolic network shows only interaction between one bacterium and one host cell type, which is far from accurately depicting the community in the gut. Nevertheless, such simplified metabolic networks represent a promising first step towards understanding the interactions between the host and gut microbiota.

## Simulation of dysbiosis and treatment using metabolic networks

As mentioned the gut microbiota is more or less stable in a healthy gut [70] and is able to recover from short term perturbations such as short-term antibiotic administration, periods of starvation, and radical changes in diet [24, 74, 75]. However, long term perturbations such as prolonged antibiotic use [76], but also changing diet can cause a shift in microbiota composition, and in turn, a shift in its metabolic products [22, 74, 77]. Alterations in microbiota composition, also known as dysbiosis, can have negative consequences on host health. Dysbiosis has been reported in patients with Alzheimer’s disease [78], Huntington’s disease [79], and Parkinson’s disease (PD) [80]. The microbiome modeling toolbox can be used to construct a metabolic network of the microbiota from the relative abundance data obtained from 16S sequencing studies of fecal samples and can be used to investigate dysbiosis [9]. Baldini et al. used the microbiome modeling toolbox to construct personalized metabolic networks of PD patients and compared differences in their metabolic output with healthy controls. The study reported 9 metabolites to have the potential to be significantly altered in PD patients compared to healthy controls including methionine and cysteinylglycine, which are part of sulfur metabolism. Furthermore the authors showed that a higher presence of *Akkermansia muciniphila*, and *Bilophila wadsworthia* in PD patients, and identified a new research target in PD research with the use of metabolic networks [81]. Hertel *et al*. confirmed these results by constructing GEMs of 31 early stage, drug naïve PD patients and 28 age matched controls from data obtained from Bedarf et al [82]. Constructing metabolic networks and simulating using FBA concluded that four microbial reactions involved in homoserine metabolism are altered in PD patients, which is consistent with measured levels in plasma of PD patients and homoserine, the precursor of methionine, is part of sulfur metabolism. Furthermore, the authors showed an increase in two bacterial species both involved in sulfur metabolism, *B. wadsworthia* and *A. muciniphila* [83]. Both species have been reported in PD progression [84]. The above-mentioned studies show the potential for metabolic networks and FBA in investigating causal relationships in understanding diseases. Additionally, dysbiosis can play a role in the treatment of neurodegenerative diseases. For example, the treatment of PD patients is done with levodopa (L-DOPA). However, the dosage varies widely among patients [85]. Members of the gut microbiota can convert L-DOPA. Thus, having more L-DOPA converting bacteria in the microbiota results in a higher dosage of L-DOPA among PD patients[86, 87]. On the contrary, medication used to treat brain-related diseases can have negative consequences by inducing dysbiosis in the gut. For example fluoxetine, a drug used as an antidepressant [88], causes sporulation in members of the gut microbiota. Thus, changing the metabolic output of the microbiota, which may impact host health [89]. The above-mentioned studies show an association between medication, and dysbiosis but lack the causality of dysbiosis and options to treat the dysbiosis to improve drug effectiveness. Constructing metabolic networks of a dysbiosis can give insight in the interactions of the dysbiosis. Next, effectiveness of treatment can be tested *in silico* on the microbiota of each patient individually, which will improve the process of choosing the best treatment, when prescribing medication.

Currently, dysbiosis can be treated in several ways such as a fecal matter transplantation (FMT), antibiotic use and using pro-, pre-, or psychobiotics. FMT has been used successfully in recurrent *Clostridium difficle* infection [90], but not as successful in treating IBD [91]. Interestingly, a study investigating FMT in IBD showed that patients treated with fecal material from one particular donor showed more response compared to patients treated with fecal matter from other donors [92]. This suggests that for successful use of FMT the microbial species and metabolites responsible for the beneficial effects of FMT need to be identified. However, to identify the differences between donors and understand why some donors are better than others, not only the microbiota and metabolite composition is needed but also the interaction among the bacteria as well as between the bacteria and the host, indicating the need for the construction of metabolic networks of the donors.

Another possible treatment for dysbiosis is the use of probiotics. Probiotics are defined as live microorganisms that, when administered in adequate amounts, confer a health benefit on the host [93]. Numerous studies have shown positive effects of single strains or mixture of probiotics on host health in animals or humans [94, 95, 96]. In humans, the effect of probiotic administration is mostly studied in clinical trials, whereby the effect on health is measured [97]. However, these study often show contradictory or unexpected results [98, 99, 100]. For example, Suez et al., reported that administering probiotics after antibiotic treatment slowed the recovery of the microbiota [75]. Whereas, other studies show that administering probiotics after antibiotic treatment does not impact the recovery of the microbiota [101, 102]. These contradicting results can be explained by differences in the probiotic strains used, types of antibiotics, dosage of the treatment, but also diet, medical history, initial microbiota composition and genetics of the patient [98, 103]. To address these problems there is a need to decipher the molecular mechanisms underlying the observed effects of probiotics. Furthermore, the viability of the probiotic *in vivo*, and the interaction within the microbiota and the metabolic output need to be investigated. In this respect, metabolic networks provide a useful tool to investigate the effect of probiotics on the gut microbiota. When a probiotic strain is added as a node in a metabolic network of a gut community, the metabolic flux distribution of the network will change. In other words, certain levels of bacterial metabolites produced in the network will be increased or reduced. In turn, this might have an effect on the microbiota composition presented in the network as shown in Figure 6. From these changes observed *in silico*, the effectiveness of probiotic treatment can be predicted [104]. However, to accurately predict the effect adding a new species to a community, the metabolic behavior of the new species should be known at the strain level and the metabolic capability under different environments should be known. *Bifidobacterium* species are widely used as probiotics [105]. However, the GEMs of *Bifidobacteria* in the AGORA database do not give the metabolic behavior in detail. For example the GEMs of *Bifidobacteria* in the AGORA database do not show growth on starch, whereas most *Bifidobacteria* can metabolize starch [106]. Therefore, careful curation of the GEMs in the AGORA database is needed before using a FBA approach in probiotic research. Devika et al. refined GEMs of 36 strains of *Bifidobacteria* including probiotic strains used in commercialized products. The authors compared the metabolic capabilities of the GEMs under 30 different environmental conditions. Based on metabolic capabilities in the different environments the *Bifidobacteria* could be divided into three groups. Based on the metabolic capabilities of the GEMs the authors hypothesized that the protective effect of the probiotic candidate *Bifidobacterium thermophilum* RBL67 on *Salmonella* and *Listeria* species comes from the production of SCFAs. Furthermore the authors hypothesize that *Bifidobacterium gallicum* DSM20093 and *Bifidobacterium kashiwanohense* DSM21854 can help relieve constipation via production of acetate [107], showing that FBA can be a useful tool for identifying new probiotic species.

**Figure 6:**
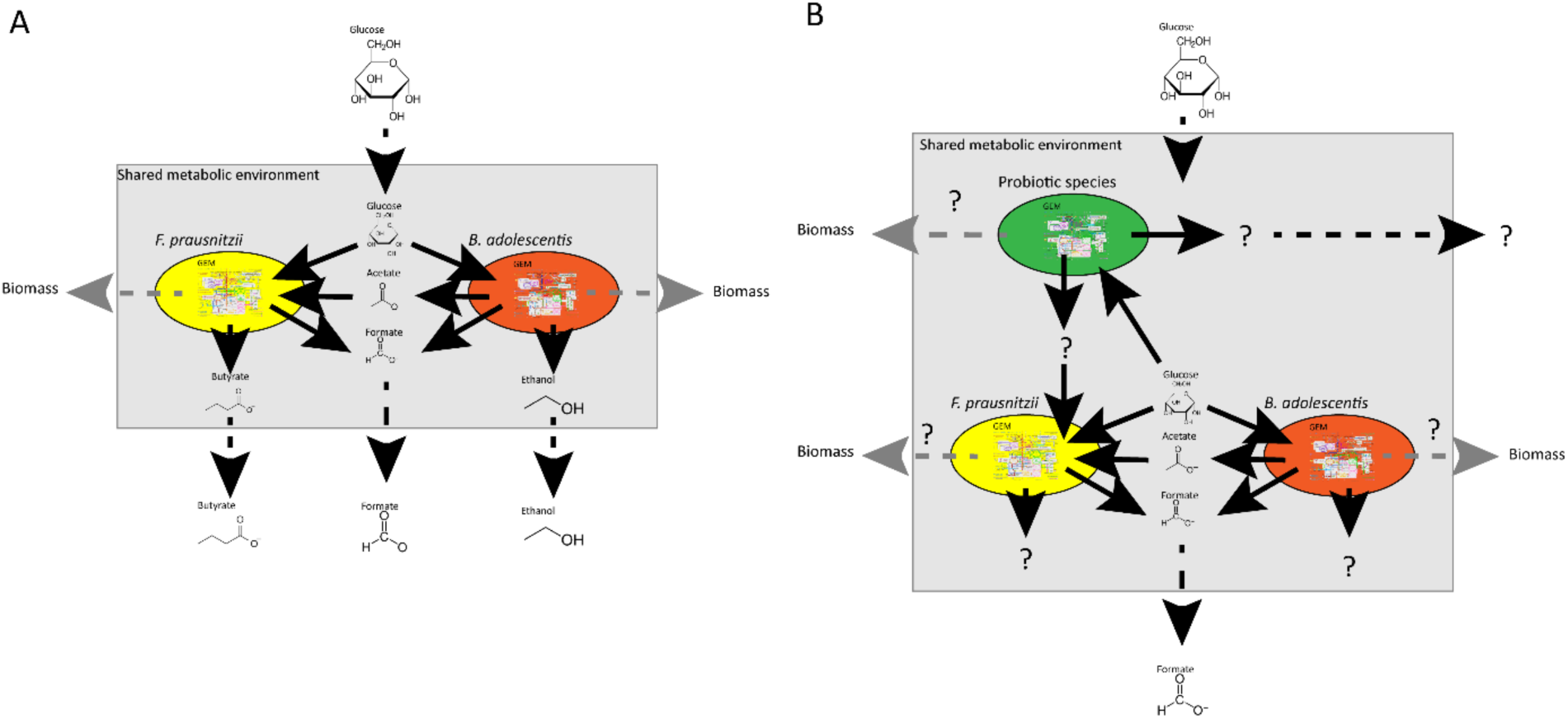
An example of the use of FBA in studying the effect of adding probiotic species in a bacterial community consisting of two gut bacteria: *F. prausnitzii* and *B. adolescentis*. (A) The community depicted in figure 5 *Adapted after El-Semman et al*. [2]. (B) Addition of another species to the shared metabolic environment will change the flux distribution of the whole system. The added species might produce a metabolite, which the other species can use, resulting in new products. Furthermore, the abundance distribution of the community might change depicted by the biomass function.

**Figure 7:**
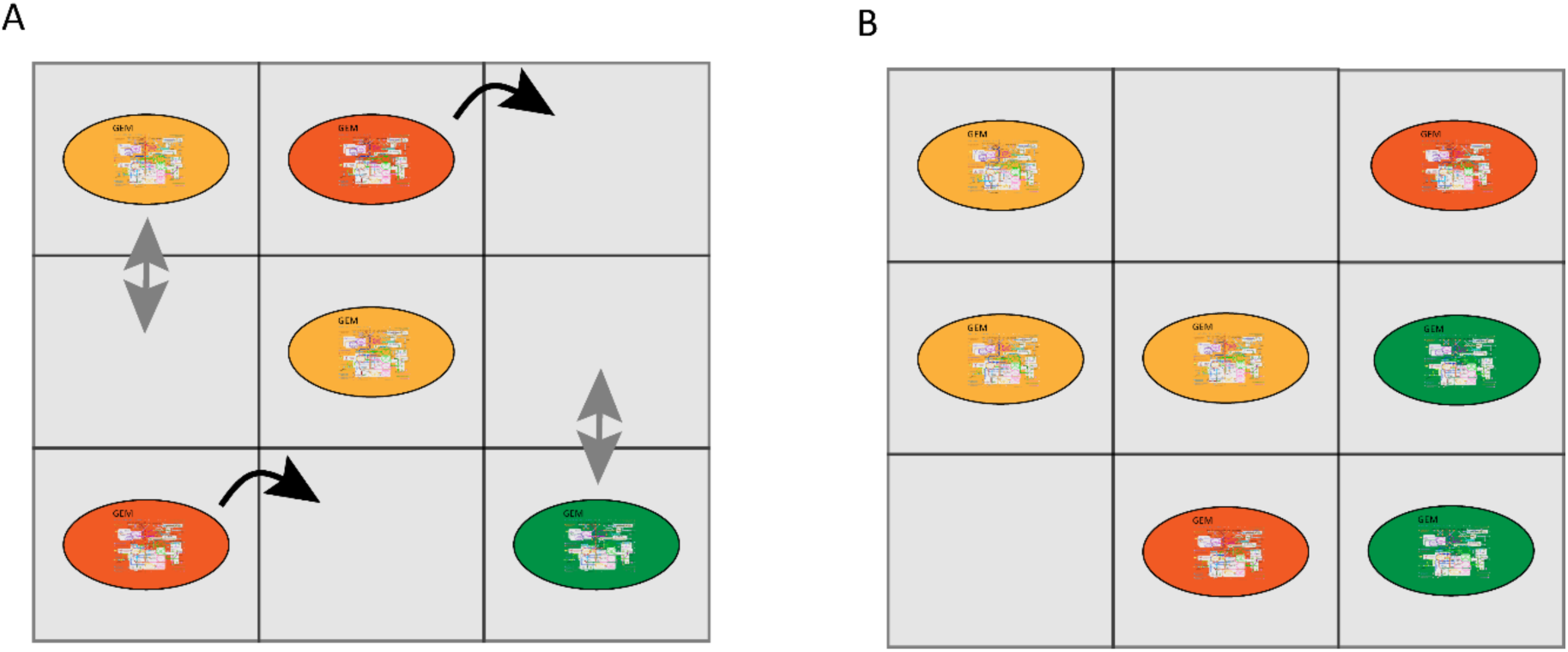
An example of the use of empty compartments in studying the gut microbiota. The metabolic compartments indicated with blue squares, are organized in a two-dimensional grid. Each bacteria has its own metabolic compartment. Exchange of metabolites can happen between the bacteria and the metabolic compartment the bacteria is in. Metabolic exchange can also happen between adjacent metabolic compartments. The metabolic compartments without a bacteria can also exchange metabolites with the adjacent compartments. In this way metabolic gradients occur. Introduction of a timestep to a grid of compartments gives the opportunity to include movement and division of bacteria. (A) The bacterial community at the start of a simulation. Movement is indicated with black, curved arrows and division is indicated with grey arrows. (B) after a timestep some cells moved and others divided, resulting in a different distribution of bacteria.

## Conclusions and Future perspectives

This review discusses the importance of combining computational (FBA-based) and experimental methods to provide novel understanding of the host-microbe, microbe-microbe interactions. Metabolic networks and FBA are promising tools to elucidate the precise interactions among bacteria, between bacteria and the host, and can be used to test the effectiveness of interventions such as probiotic treatment before clinical trials. Furthermore, metabolic networks can help point at knowledge gaps. For example, if there is a discrepancy between the outcome of model simulations, and experimental work, a more thorough understanding of the metabolic capabilities of the organism of interest is needed, which leads to new hypotheses and more experiments. Next, metabolic networks can elucidate causality in the microbiota research, because unlike in experimental work, numerous alterations can be implemented in metabolic networks [108]. Currently, metabolic networks of the gut microbiota are mainly based on databases from samples collected from western individuals [109]. To make sure that metabolic networks can be used globally, more global sampling is needed. Furthermore, to ensure the accuracy of the predicted output of the metabolic networks, sufficient information of concentrations of microbial metabolites in the gut, microbial composition, metabolic potential, and interactions is required to be implemented in the metabolic network [107, 110]. Furthermore, metabolic network models are mathematical descriptions of reality. To make sure the metabolic networks depict reality accurately, the outcome of simulating metabolic networks should be experimentally validated. However, the majority of the gut bacteria are not yet culturable [111]. Therefore, GEMs of those bacteria cannot be tested for accuracy. Furthermore, investigating interactions between bacteria can be challenging. Nevertheless, research is performed to identify, culture and characterize new species from the gut microbiota [112]. Currently, new *in vitro* methods to experimentally validate metabolic networks become available such as SHIME, HuMiX and others [108, 113, 114]. Medlock et al. combined metabolic networks with co-culturing to infer cross feeding between members of the Altered Schaedler Flora (ASF) and validated that this metabolic interaction leads to a growth benefit for the bacteria [115]. An example of the use of computational methods to benefit human health is a study performed by Zeevi et al. where a machine learning algorithm advised a personalized diet based on, blood glucose levels and microbiota composition, to lower the glycemic response after a meal. Indeed, when participants followed the advised diet, a lower glycemic response was observed [116], indicating that algorithms can predict the influence of diet on the host if data on microbial composition and host parameters are gathered. However, since this approach was based on machine learning, not an FBA, the exact metabolic interactions were not elucidated in the study. Combining computational and experimental progress can help to better understand the intricate relationship between the microbiota and the host. Over time, this understanding can be used to design targeted, and personalized approaches to alter the microbiota in a way that it benefits the health of the host.

## List of abbreviations

COBRA: Constraint-based reconstruction and analysis
FBA: Flux Balance Analysis
OF: Objective function
SCFA: Short chain fatty acid
DFBA: Dynamic flux balance analysis
ODE: Ordinary differential equations
SOA: Static optimization approach
GEM: Genomic-scale metabolic model
GENRE: Genomic-scale metabolic reconstruction
LCMS: Liquid chromatography mass spectrometry
RP: Reverse phase
HILIC: Hydrophilic interaction liquid chromatography
PD: Parkinson’s disease
L-DOPA: Levodopa
FMT: Fecal matter transplantation

## Declarations

### Ethics approval and consent to participate

Not applicable

### Consent for publication

Not applicable

### Availability of data and material

The manuscript does not include any supporting data.

## Authors’ contributions

Conceptualization, J.J., S.E.A.; Writing-Original draft, J.J.; Writing-Review & Editing, S.E.A; Funding Acquisition, S.E.A.

## Acknowledgements

We thank Dr. Andreas Milias-Argeitis of Department of Molecular Systems Biology, University of Groningen, The Netherlands for critical reading of our manuscript.

